# PARAQUAT TOLERANCE3 is an E3 ligase and acts as a negative regulator of oxidative stress response

**DOI:** 10.1101/040543

**Authors:** Chao Luo, Xiao-Teng Cai, Jin Du, Tao-Lan Zhao, Peng-Fei Wang, Ping-Xia Zhao, Rui Liu, Qi Xie, Xiao-Feng Cao, Cheng-Bin Xiang

## Abstract

Oxidative damage could be caused in plant cells when biotic and abiotic stresses are imposed. While the response to oxidative stress is well studied, little is known about how the activated response is switched off when oxidative stress is diminished. By studying Arabidopsis mutant *paraquat tolerance3*, we identified the genetic locus *PARAQUAT TOLERANCE3* (*PQT3*) as a major negative regulator of oxidative stress tolerance. *PQT3*, encoding an E3 ligase, is rapidly down-regulated by oxidative stress. PQT3 has E3 ubiquitin ligase activity in ubiquitination assay. Subsequently, we identified PRMT4b as a PQT3-interacting protein. By histone methylation, PRMT4b may regulate the expression of *APX1* and *GPX1*, encoding two key enzymes against oxidative stress. Moreover, PQT3 is able to recognize PRMT4b for targeted degradation via 26S proteasome. Therefore, we have identified PQT3 as an E3 ligase that acts as a negative regulator of activated response to oxidative stress.

---

Sessile plants cannot avoid harsh living conditions such as drought, salinity, cold and hot temperature. These stresses induce damages to plants in different degrees. In most cases, they alter the normal cell homeostasis and increase the generation of reactive oxygen species (ROS) (Miller et al., 2010). Exposing plants to excessive light also causes ROS accumulation, serious damage to the photosynthetic system, and photo-inhibition (Mittler, 2002; Caverzan et al., 2012). Excessive ROS production results in oxidative stress which could destroy biological membranes and macromolecules, accelerate cell senescence, induce irreversible damages to cells and even lead to cell death (Gill and Tuteja, 2010). Under normal circumstances, ROS can be controlled at a low level in the organelles where ROS are produced, such as chloroplasts, mitochondria, and peroxisomes. However, the level of ROS is sharply increased under stress conditions (Moller, 2001; Mittler et al., 2004; del Rio et al., 2006; Leshem et al., 2006; Corpas et al., 2008; Miller et al., 2008; Van Breusegem et al., 2008; Miller et al., 2010). The production and elimination of ROS are dynamically balanced in plant cells (Apel and Hirt, 2004). Two protection systems, enzymatic and non-enzymatic, have evolved to scavenge ROS and protect plant cells from oxidative stress. Enzymatic system mainly includes the superoxide dismutase (SOD), ascorbate peroxidase (APX), catalase (CAT), glutathione peroxidase (GPX) and peroxiredoxin (PrxR), while non-enzymatic system mainly contains the ascorbic acid (AsA), alpha tocopherol (VE), and glutathione (GSH) (Takahashi and Asada, 1988; Apel and Hirt, 2004; Mittler et al., 2004). ROS are also important signal molecules that mediate the responses to pathogens, environmental stress, and stimuli in various developmental processes (Mittler et al., 2004; Torres and Dangl, 2005). The coordination mode of different networks of ROS elimination is very complex in Arabidopsis (Mittler et al., 2004).

The activation of oxidative response involves many layers of regulations (Mittler, 2002; Mittler et al., 2004). Little is known about the regulation by histone methylation of the genes involved in oxidative stress response. Histone methylation plays important roles in the plant development and growth as well as in some stress responses (Niu et al., 2007; Niu et al., 2008; Hong et al., 2010; Liu et al., 2010a; Yuan et al., 2013). The methylation marks are written on lysines or arginines respectively by histone lysine methyltransferases (HKMTs) and protein arginine methyltransferases (PRMTs). Nine PRMTs are found in the Arabidopsis genome (Niu et al., 2007). Two different types of PRMTs catalyze asymmetric di-methylation (ADMA) and symmetric di-methylation (SDMA) on the Arg residues, respectively (Scorilas et al., 2000; Liu et al., 2010a). A pair of PROTEIN ARGININE METHYLTRANSFERASE4 (PRMT4) homologs, AtPRMT4a and AtPRMT4b, is required for the asymmetrical di-methylation of Arg-2, Arg-17, and Arg-26 in histone H3 (Niu et al., 2007). Protein arginine methylation plays essential roles in diverse biological processes, such as RNA processing and transcriptional regulation (Scorilas et al., 2000).

Oxidative stress could be perceived by multiple mechanisms, including sensor or cellular receptor. The perceiption by receptors results in the activation of Ca^2+^ – calmodulin and mitogen-activated protein kinase (MAPK) cascade signaling transduction pathway. The activation or suppression of different transcription factors regulates a variety of defense pathway subsequently, such as ROS-scavenging, heat-shock proteins (HSPs), and photosynthesis (Mittler, 2002; Apel and Hirt, 2004; Mittler et al., 2004). While much attention has been paid to how plants respond to oxidative stress, we know little about how plants switch off the activated responses when stress is diminished. A common regulatory mechanism is to control the protein level of the stress responsive factors. The most studied mechanism of protein degradation is the ubiquitin/26S proteasome system (Santner and Estelle, 2010). The ubiquitin/26S proteasome pathway has been implicated in diverse aspects of eukaryotic cell regulation because of its ability to rapidly remove specific intracellular proteins (Moon et al., 2004; Dreher and Callis, 2007; Santner and Estelle, 2010). In this system, ubiquitin composed of 76 amino acids, is attached to the target protein under the action of three different enzymes (Dreher and Callis, 2007). The target proteins with ubiquitin have different fates. One kind of the ubiquitinated proteins can be recognized and degraded by the 26S proteasome. Another type of labeled proteins, which are folded in wrong ways, can be recognized by their related enzymes to achieve the correct protein structures. The ubiquitin system may also directly affect protein activity and intracellular localization. It can also identify and modify many intracellular proteins, such as proteins involved in signal transduction, transcription factors, and receptors on cell surface, to participate in the regulation of physiological processes (Mukhopadhyay and Riezman, 2007).

In the ubiquitin degradation process, E3 played a crucial role. E3 is responsible for specific recognition of substrate protein and accurate positioning of the binding site between substrate protein and ubiquitin (Yee and Goring, 2009). In *Arabidopsis thaliana*, approximately 1400 genes encode components of the ubiquitin/26S proteasome pathway (Smalle and Vierstra, 2004). Approximately 90% of these genes encode subunits of the E3 ubiquitin ligases (Moon et al., 2004). Based on the domain interacted with E2 (ubiquitin conjugase) during the ubiquitin conjugation cascade, the large and diverse family of plant E3 ubiquitin ligases can be divided into HECT domain- and RING/U-box domain-containing E3 ligases (Santner and Estelle, 2010). The HECT family is relatively small compared with the RING domain-containing family that includes several hundreds of proteins and can be further divided into single subunit RING/U-box E3 ligases and multi subunit RING E3 ligases (Pickart, 2001; Moon et al., 2004). The large number of E3 ubiquitin ligases in higher plants indicates their important regulatory roles in diverse biological processes (Mazzucotelli et al., 2006).

Few E3 ligases have been identified to turn off the activated stress responses and the function of an enormous number of E3 ligases still remains to be identified. Here we report a novel negative regulator of oxidative stress response by PARAQUAT TOLERANCE3 (PQT3) in Arabidopsis. We isolated a paraquat tolerant mutant, ***p****ara****q****uat* ***t****olerance3* (*pqt3*), and cloned the gene *PQT3* that encodes an E3 ligase containing RING/U-box domain. The expression of *APX1* and *GPX1* was up-regulated in *pqt3*, while *PQT3* was down regulated by oxidative stress. PQT3 was able to interact with PRMT4b. PRMT4b may catalyze histone methylation on *APX1* and *GPX1* chromatin and up-regulate their expressions, therefore protect plants from oxidative stress. When oxidative stress is diminished, PQT3 level increases and acts as E3 ubiquitin ligase to specifically target PRMT4b for degradation. Based on our results, PQT3 is a negative regulator that turns off the activated response of oxidative stress.

## Results

### Loss of *At4g17410* confers the paraquat tolerance of *pqt3*

The mutant *pqt3* was isolated from an activation-tagging library as described (Xi et al., 2012). This library containing approximately 55, 000 independent lines was screened for mutants with enhanced tolerance to different stresses (Yu et al., 2008; Wu et al., 2010). To isolate tolerant mutant to oxidative stress, we germinate seeds on MS medium with 2 μM paraquat, green seedlings that could continued to grow were rescued as putative mutants with improved paraquat tolerance and named *paraquat tolerance* (*pqt*) mutants. The *pqt3*, as one mutant of them, was further characterized and marked as *pqt3-1*. The enhanced oxidative tolerance of *pqt3-1* mutant was confirmed by germinating seeds on MS medium containing 0 or 2 μM of paraquat. In presence of paraquat, more than 60% *pqt3-1* seeds germinated with green cotyledons but only 2% wild type seeds germinated, while all seeds of both wild type and *pqt3-1* germinated on MS medium without paraquat (Figures 1A and 1B). Genetic analysis showed that the mutation was recessive. All F1 backcross offsprings (*pqt3-1* x wild type) were paraquat sensitive and F2 selfing population showed typical 3:1 segregation ratio (sensitive: resistant; 85:27, χ^2^ = 0.0476). The result suggested that *pqt3-1* mutant may have a more efficient mechanism of ROS scavenge, which was caused by loss-of-function mutation in a single nuclear gene *PQT3* (*At4g17410*).

**Figure 1.**
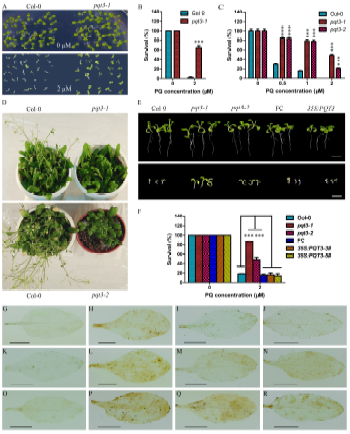
Phenotype of *pqt3-1* and *pqt3-2* mutants. **(A)** Confirmation of paraquat (PQ) tolerant phenotype. 7-day-old seedlings of wild type and *pqt3* mutant were grown on MS medium supplemented with 0 or 2 μM paraquat. **(B)** Survival ratio of wild type and *pqt3* mutant grown in **(A)**. Values are mean ±SD (n= 30 plants, ***P < 0.001). Asterisks indicate Student’s t-test significant differences. **(C)** Multiple mutant alleles analysis of paraquat tolerance for the *At4g17410* locus. Survival ratio of wild type, *pqt3-1* and *pqt3-2* (Salk_065409) grown in the 0, 0.5, 1, and 2 μM paraquat medium was counted. Values are mean ±SD (n= 30 plants, ***P < 0.001). Asterisks indicate Student’s t-test significant differences. **(D)** Late-flowering phenotype of *pqt3-1* and *pqt3-2* compared with wild type. Plants were grown under long day photoperiod (16 h light and 8 h dark). **(E)** The phenotype of 7-day-old *pqt3-1*, wild type, function complementation (FC), *pqt3-2*, and *35Spro:PQT3* seedlings under 0 μM (the top picture) or 2 μM (the bottom picture) paraquat treatment. Bar= 0.5 cm. **(F)** The survival ratio of *pqt3-1*, wild type, FC, *pqt3-2*, and *35Spro:PQT3* seeds germinated and grown on MS medium containing 0 μM or 2 μM paraquat for 7 days was counted. Values are mean ±SD (n= 30 plants, ***P < 0.001). Asterisks indicate Student’s t-test significant differences. **(G to R)** DAB staining. The leaves of wild type, *pqt3-1*, and *pqt3-2* were treated without **(G to J)** or with 6 μM paraquat for 12 **(K to N)** or 24 h **(O to R)**. 10 mM Na_2_HPO_4_ was used as the negative control to stain the leaves of wild type treated without **(G)** or with 6 μM paraquat for 12 **(K)** or 24 h **(O)**. Without PQ treatment, the leaves of wild type **(H)**, *pqt3-1* **(I)**, and *pqt3-2* **(J)** were stained using DAB staining solution. After 12 h paraquat treatment, DAB staining solution was used to stain the leaves of wild type **(L)**, *pqt3-1* **(M)**, and *pqt3-2* **(N)**. The leaves of wild type **(P)**, *pqt3-1* **(Q)**, and *pqt3-2* **(R)** were stained by DAB staining solution after 24 h paraquat treatment. Bar= 0.5 cm.

In *pqt3-1* mutant, a single T-DNA insertion was located in the fourth intron of *At4g17410* (Supplemental Figure 1A). The exact integration site of the T-DNA right border was 803bp downstream of the ATG initiation codon of *At4g17410*. As a result, the expression of *At4g17410* was completely disrupted as confirmed by RT-PCR analysis (Supplemental Figures 1B and 1D). The expressions of its neighboring genes, *At4g17390* and *At4g17420*, were not affected (Supplemental Figure. 1B).

The *At4g17410* locus consists of 13 exons and 12 introns. The open reading frame is predicted to encode a polypeptide composed of 827 amino acids with an estimated molecular mass of 91kD. Based on the conserved RING/U-box domain, this protein is predicted as an E3 ligase.

To further determine whether the loss of *At4g17410* resulted in the enhanced oxidative tolerance of *pqt3-1* mutant, we used another allele of *pqt3*, the T-DNA insertion mutant Salk_065409, which was ordered from Arabidopsis Biological Resource Center ( ABRC) and its T-DNA insertion was confirmed by RT-PCR (Supplemental Figures 1A, 1C and 1D). As the first identified *pqt3* mutant was named as *pqt3-1*, the Salk_065409 was marked as *pqt3-2*. The *pqt3-2* mutant showed similar enhanced oxidative tolerance to paraquat and had high survival ratio under different concentrations of paraquat treatment as *pqt3-1* did (Figure 1C). The survival ratio of *pqt3-1* and *pqt3-2* were 50% and 20%, respectively, under 2 μM paraquat treatment, while none of the wild type seedlings survived under the same condition. In addition, both *pqt3-1* and *pqt3-2* showed a late-flowering phenotype (Figure 1D). To confirm further, we generated functional complementation (FC) lines and *35Spro:PQT3* overexpression lines (Supplemental Figures. 1E and 1F). FC lines and *35Spro:PQT3* lines showed similar if not higher paraquat sensitivity to wild type under 2 μM paraquat treatment while the *pqt3-1* and the *pqt3-2* mutants displayed enhanced paraquat tolerance (Figures 1E and 1F). These results indicate that PQT3 is a negative regulator of oxidative stress tolerance and is responsible for the phenotype of *pqt3* mutants.

After 6 μM paraquat treatment for 12 or 24 hours, the result of DAB staining showed that the brown precipitate on the leaves of the wild type was more than that of *pqt3* mutants (Figures 1G to 1R). As several stresses could cause oxidative damage to plants, the sensitivity of *pqt3* mutants to other environmental stresses was analyzed subsequently. The result indicated that *pqt3* mutants have enhanced tolerance to CdCl_2_, mannitol, NaCl, and drought stress (Supplemental Figure 2).

### The expression pattern and protein localization of *PQT3*

To investigate the spatiotemporal pattern of *PQT3* expression, we generated *PQT3pro:GUS* reporter lines. GUS staining results showed that *PQT3* was expressed in both shoot and root tissues under normal condition (Figures 2A to 2G). GUS expression was detected in the root tissues at all developmental stages we analyzed (Figures 2A to 2C). For the 1-week-old seedlings, strong GUS staining was observed in the cotyledons, hypocotyls and root tissues (Figure. 2A). For the 3-week-old seedlings, strong GUS staining was also detected in cotyledons, young leaves, and root tissues, but weakly stained in older leaves (Figure. 2B). In 7-week-old adult plants, GUS expression was detected in rosette leaves, cauline leaves, the tip and basal junction of siliques, and was significantly higher in the flower petals, stamens and stigma of pistil (Figures. 2D to 2G).

**Figure 2.**
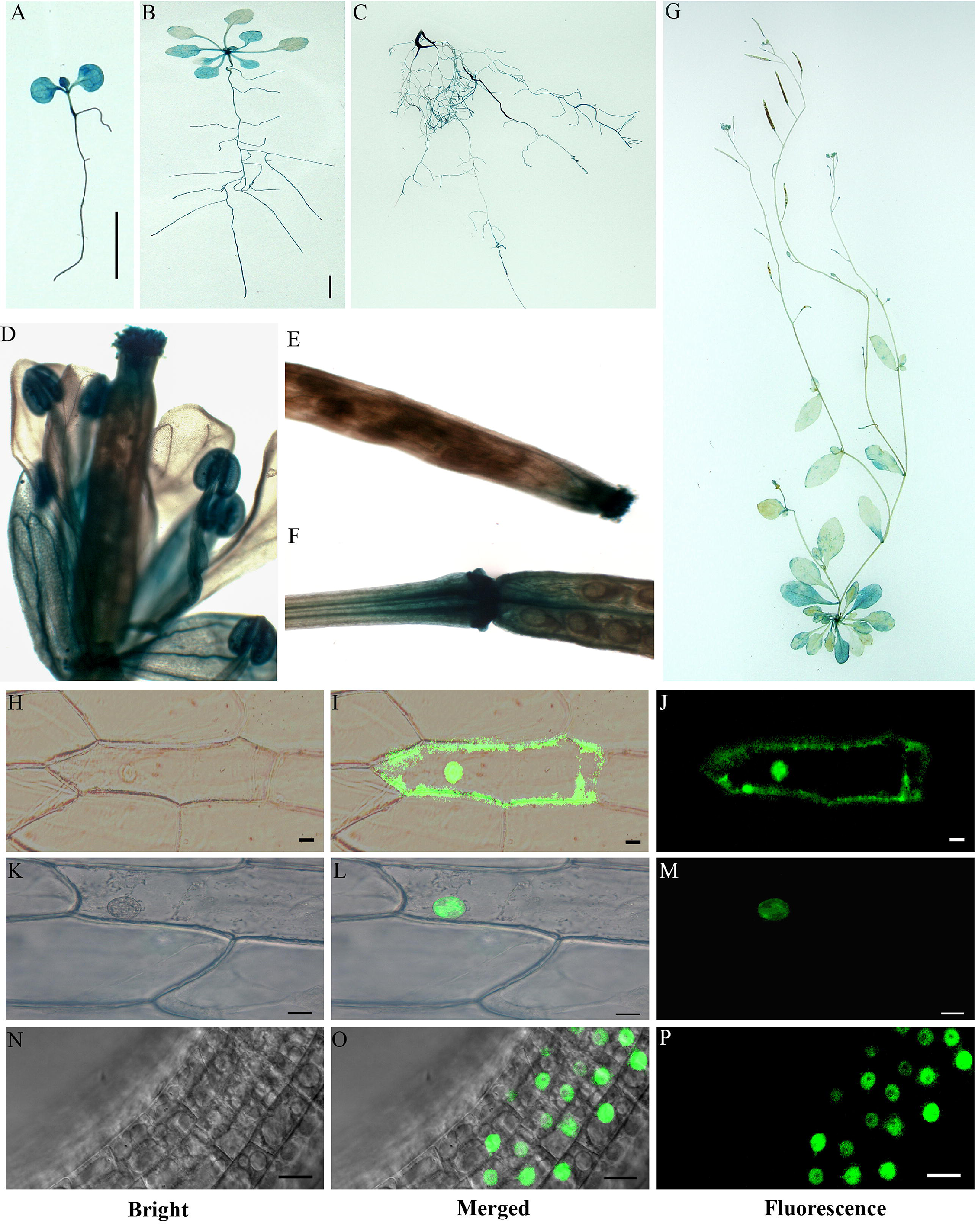
Expression pattern and subcellular localization of PQT3. **(A)** GUS expression in 1-week-old seedling. Transgenic Arabidopsis plants expressing *PQT3pro:GUS* were generated and analyzed for GUS expression. Bar= 0.5 cm. **(B)** GUS expression in 3-week-old seedling. Bar= 0.5 cm. **(C)** GUS expression in root tissue of 7-week-old plant. **(D)** GUS expression in flower tissue. **(E and F)** GUS expression in silique tip **(E)** and junction **(F). (G)** GUS expression in 7-week-old adult Arabidopsis. **(H to J)** *35Spro:GFP* was transiently expressed in onion epidermal cells as control. The GFP can be observed in both plasma membrane and nucleus. **(K to M)** Nucleus localization of the PQT3-GFP fusion protein in onion epidermal cell was observed. Bar= 20 μm. **(N to P)** The nucleus localization of PQT3-GFP fusion protein in the root tissue of stable transgenic seedlings expressing *PQT3pro:PQT3-GFP*. Bar= 20 μm.

PQT3 has two predicted nuclear localization signals (NLSs) in the carboxyl terminus (Supplemental Figure 3), implicating its nuclear localization. To confirm this, the *35Spro:PQT3-GFP* construct was made and transiently expressed in onion epidermal cells. The *35Spro:GFP* construct was used as control (Figures 2H to 2J). PQT3-GFP signal was indeed detected in the nucleus (Figures 2K to 2M).

For further confirmation, we transformed the *PQT3pro: PQT3-GFP* fusion construct into the Arabidopsis and obtained transgenic plants. Fluorescent microscopy results showed that GFP signal was accumulated in the nucleus of root cells (Figures. 2N to 2P), which is in agreement with the presence of two NLSs of the PQT3.

### PQT3 is rapidly down regulated by paraquat treatment and other stress conditions

Interestingly, the expression of *PQT3* was quickly down-regulated by 6 μM paraquat treatment and maintained at a low level as long as the paraquat treatment was applied (Figure 3A). This result is supportive for our previous opinion that PQT3 is a negative regulator of plant oxidative tolerance. By extropolation, the expression of *PQT3* could be down-regulated by other stress conditions. Indeed, our results showed that the expression of *PQT3* was down-regulated by H_2_O_2_ (Figure 3B), mannitol (Figure 3C), drought (Figure 3D), CdCl_2_ (Figure 3E), and NaCl treatment (Figure 3F) at the indicated time points. Among these stresses, CdCl_2_ treatment led to the most significant reduction of *PQT3* expression (Figure 3E). The *PQT3pro:GUS* pattern was observed under differrent stresses subsequently (Figures 3G to 3DD). Compared with transgenic seedlings under normol conditions (Figures 3G, 3M, 3S and 3Y), GUS staining was weaker in the seedlings under paraquat (Figures 3H, 3N, 3T and 3Z), H_2_O_2_ (Figures 3I, 3O, 3U and 3AA), mannitol (Figures 3J, 3P, 3V and 3BB), CdCl_2_ (Figures 3K, 3Q, 3W and 3CC), and NaCl treatment (Figures 3L, 3R, 3X and 3DD). The changed *PQT3pro:GUS* pattern was consistent with the altered expression of *PQT3* detected by quantitative RT-PCR under differrent stresses.

**Figure 3.**
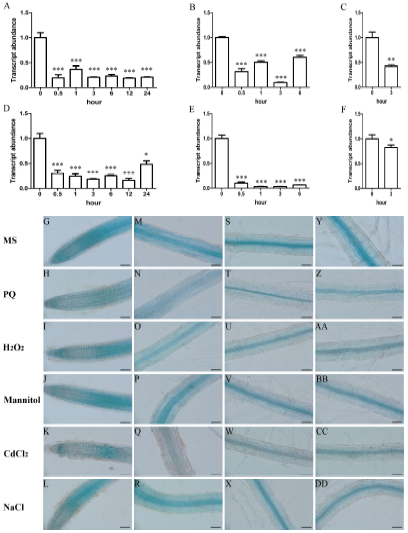
The expression of *PQT3* was down-regulated by various oxidative stress conditions. **(A)** The transcript level of *PQT3* was down-regulated by paraquat treatment. 1-week-old seedlings were treated with 6 μM paraquat for the indicated times before RNA extraction for quantitative RT-PCR analysis. Values are mean ±SD (n=3 experiments, ***P < 0.001). Asterisks indicate Student’s t-test significant differences. **(B to F)** The expression of *PQT3* was down-regulated by other stress conditions. 1-week-old seedlings were treated by 10 mM H_2_O_2_ **(B)**, 200 mM mannitol **(C)**, drought **(D)**, 200 mM CdCl_2_ **(E)**, and 150 mM NaCl **(F)** for the indicated times before RNA extraction for quantitative RT-PCR analysis. Values are mean ±SD (n=3 experiments, *P < 0.05, **P < 0.01, ***P < 0.001). Asterisks indicate Student’s t-test significant differences. **(G to DD)** GUS staining of 7-day-old transgenic lines contained *PQT3pro:GUS* without or with 6 μM paraquat, 10 mM H_2_O_2_, 200 mM mannitol, 200 mM CdCl_2_, 150 mM NaCl treatment for 3 h. GUS expressions were significantly reduced in primary root tip **(G to L)**, root elongation zone **(M to R)**, root maturation zone **(S to X)**, root zone with LRP **(Y to DD)** under stress conditions. Bar= 50 μm.

### *APX1* and *GPX1* were up-regulated in *pqt3* mutant

Enzymatic protection systems were very important for ROS elimination. The transcript levels of ascorbate peroxidase (APX), catalase (CAT), glutathione peroxidase (GPX), cytosolic Cu/Zn SOD (CSD1), plastidic Cu/Zn SOD (CSD2), FeSOD (FSD), atypical Cys-His rich thioredoxin (ACHT), glutaredoxin C (GRXC), 2-Cys peroxiredoxin B (2CPB), peroxiredoxin Q (PRXQ), and mitochondrial MnSOD (MSD) were analyzed by quantitative RT-PCR in *pqt3* and wild type. The results show that transcript levels of *APX1* and *GPX1* were up-regulated in *pqt3* under normal conditions compared with that in the wild type (Figures 4A to 4J). The elevated transcript levels of *APX1* and *GPX1* may contribute to the enhanced oxidative stress tolerance. The enzyme activity of APX and GPX in wild type and *pqt3* mutant was also detected. The *pqt3* mutant had higher enzyme activity of APX and GPX compared with wild type (Figures 4K and 4L).

**Figure 4.**
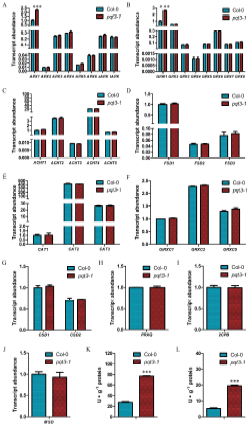
The analysis of antioxidant enzyme in wild type and *pqt3* mutant. **(A to J)** Quantitative RT-PCR analysis of transcript levels of antioxidant enzyme genes. RNA samples were isolated from 7-day-old wild type and *pqt3* seedlings for quantitative RT-PCR analysis. The transcript levels of APX **(A)**, GPX **(B)**, ACHT **(C)**, FSD **(D)**, CAT **(E)**, GRXC **(F)**, CSD **(G)**, PRXQ **(H)**, 2CPB **(I)** and MSD **(J)** were analyzed. Values are mean ±SD (n=3 experiments, ***P < 0.001). Asterisks indicate Student’s t-test significant differences. **(K and L)** Enzyme activity of APX **(K)** and GPX **(L)** in wild type and *pqt3* mutant. The *pqt3* mutant has higher enzyme activity of APX and GPX than wild type. Values are mean ±SD (n=3 experiments, ***P < 0.001). Asterisks indicate Student’s t-test significant differences.

### PQT3 possesses E3 ligase activity

Not all the proteins with the predicted RING domain function as an ubiquitin ligase (Deshaies and Joazeiro, 2009). The E3 activity of PQT3 was determined via self-ubiquitination system. The full-length PQT3 protein (GST-PQT3) showed the E3 ubiquitin ligase activity (Figure 5). The ubiquitinated bands of PQT3 were detected by western blotting in the presence of E1 (from wheat), E2 (UBCh5b, from human), and 6×His-tagged ubiquitin (UBQ14, from Arabidopsis). Self-ubiquitination of PQT3 was not detected in the absence of any of the essential reaction components (Figure. 5). The result demonstrates that PQT3 has the E3 ubiquitin ligase activity.

**Figure 5.**
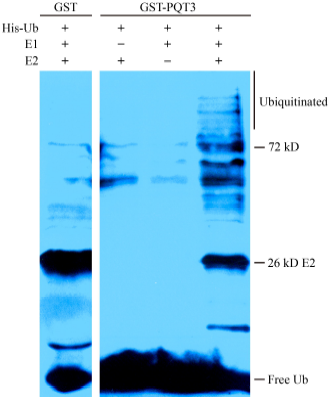
PQT3 has E3 ubiquitin ligase activity. GST-PQT3 was expressed in *E. coli* and purified. Nickel-HRP was used to detect His-tagged ubiquitin. The ubiquitination activity of GST-PQT3 was observed only in the presence of E1 (from wheat), E2 (UBCh5b, from human), and 6×His-tagged ubiquitin (Ub). GST was used as negative control. The numbers on the right indicate the molecular masses of marker proteins.

### PQT3 interacts with PRMT4b

To study the molecular mechanisms that underlie the enhanced stress tolerance of *pqt3*, we screened cDNA library for potential candidate target proteins of PQT3 using yeast-two-hybrid (Y2H). Several proteins were isolated from the screen. Among these candidate interactors, PRMT4b, a member of arginine methyl transferase family, was frequently presented. To reveal the domain of PQT3 responsible for the interaction with PRMT4b, the PQT3 protein was divided into four parts: N-terminal DWNN, zfCCHC, U-box (RING finger), and C-terminal section containing the NLS1 and NLS2 domains, based on the predicted domains of PQT3 protein (Supplemental Figure 3). Full-length PQT3 and four protein sections were used for Y2H assay as baits. Full-length protein and the C-terminus containing the NLS1 and NLS2 domains of PQT3 were able to interact with PRMT4b in Y2H assays (Figures 6A to 6C). The interaction between PQT3 and PRMT4b was confirmed by colonies that grew on the SC-Leu-Trp-His plate with 50 mM 3-amino-1,2,4-triazole (3-AT) and displayed the blue color in X-gal assay (Figures 6B and 6C). PRMT4a is not a potential target of PQT3 in previous Y2H library screening. The Y2H assay was also performed to further study the potential interaction between PQT3 and PRMT4a, since *PRMT4a* is a close related gene of *PRMT4b*. The result showed that the PQT3 can not interact with PRMT4a (Supplemental Figure 4).

**Figure 6.**
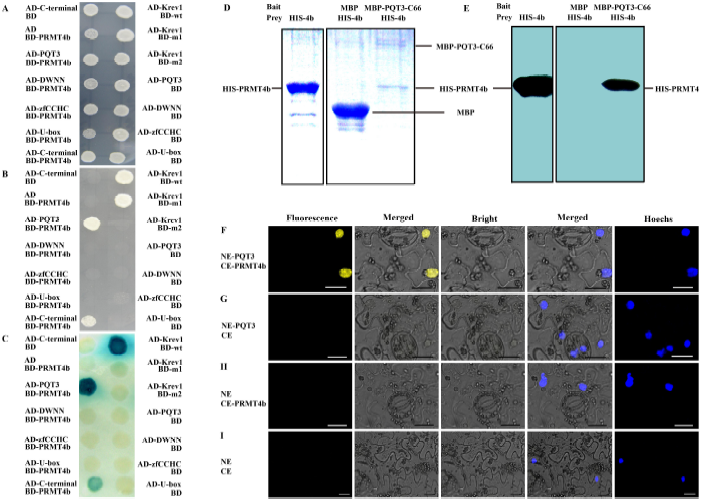
PQT3 interacts with PRMT4b. **(A to C)** Y2H assay. PQT3 and its four protein sections were used as the bait. PRMT4b was used as the prey. Krev1/RalGDS-wt act as strong positive control and Krev1/RalGDS-m1 act as week positive control. Krev1/RalGDS-m2 was used for negative control. The yeast harboring various constructs was grown on SC-Leu-Trp medium **(A)**. The yeast was transferred to SC-Leu-Trp-His medium with 50 mM 3-AT **(B)** or used for X-gal staining **(C)**. **(D and E)** The pull-down assay between PQT3 and PRMT4b. His-PRMT4b was incubated with amylose resin bound with recombinant MBP-PQT3-C66 protein. Pulled-down protein complex was detected by SDS-PAGE **(D)** and western blot using anti-His antibody **(E)**. MBP protein was used as a negative control. **(F to I)** BiFC assay. Different plasmid combinations were expressed in epidermal cells of *N. benthamiana* leaves. Yellow fluorescence (YFP) was observed in epidermal cell expressing both NE-PQT3 (the N-terminal part of YFP fused with PQT3) and CE-PRMT4b (the C-terminal part of YFP fused with PRMT4b) **(F)**. No fluorescence was observed from the negative controls: NE-PQT3 / CE **(G)**, NE / CE-PRMT4 **(H)**, and NE / CE **(I)**. The nuclei were stained by Hoechst and the fluorescence was detected by confocal. NE indicates pSPYNE vector and CE indicates pSPYCE vector.

Protein pull-down assay was performed to confirm the interaction between PQT3 and PRMT4b *in vitro*. MBP-PQT3-C66 protein containing the NLS domain (Supplemental Figure 3) and His-PRMT4b protein were expressed in *E. coli* and purified subsequently. His-PRMT4b was incubated with amylose resin bound with recombinant MBP-PQT3-C66 protein. Pulled-down protein complex was detected by SDS-PAGE (Figure 6D) and western blotting using anti-His antibody (Figure 6E). The pull-down result clearly shows that PQT3 interacts with PRMT4b *in vitro*.

To determine whether the interaction also occur *in vivo*, we used the bimolecular fluorescence complementation (BiFC) system. Full-length *PQT3* cDNA was fused to the N-terminal region of the yellow fluorescent protein (YFP), while full-length *PRMT4b* cDNA was fused to the C-terminal region of YFP. The empty plasmids were used as negative controls. Different plasmid combinations were co-infiltrated into epidermal cell of *N. benthamiana* leaves. The yellow fluorescence was observed in epidermal cell contained both NE-PQT3 (the N-terminal part of YFP fused with PQT3) and CE-PRMT4b (the C-terminal part of YFP fused with PRMT4b) (Figure 6F). No fluorescence was observed from the negative controls (NE-PQT3/CE, NE/CE-PRMT4b and NE/CE) (Figures 6G, 6H and 6I). The nuclei were stained by Hoechst and detected by confocal subsequently. These results indicate that PQT3 can interact with PRMT4b in the nucleus of plant cell.

### PRMT4b is an ubiquitination substrate of PQT3

To determine whether PRMT4b is a substrate recognized by PQT3 being as an E3-ligase, we resorted to the *in planta* ubiquitination assay (Liu et al., 2010b). Leaf infiltration was conducted via *Agrobacterium tumefaciens* strains containing different combination of constructs. The infiltrated parts of *N. benthamiana* leaves were harvested. Total protein was extracted and detected via western blotting with anti-HA antibody. A smear of bands, which were larger than the size of HA-PRMT4b and showed the features of ubiquitinated form of the PRMT4b proteins, could be examined by anti-HA antibody in the samples co-infiltrated with PQT3 and HA-PRMT4b (Figure 7A). The cell lysates were immunoprecipitated with anti-HA antibody subsequently. Immunoprecipitated samples were detected via western blotting with anti-ubiquitin antibody. In the PQT3-PRMT4b co-infiltration sample, these high molecular size bands could also be detected by anti-ubiquitin antibody (Figure 7B). These results indicated that these high molecular size bands were ubiquitinated forms of PRMT4b. PQT3 protein could ubiquitinate the PRMT4b protein in tobacco. The decline of PRMT4b protein was also found in the samples co-infiltrated with PQT3 and HA-PRMT4b (Figure 7A).

**Figure 7.**
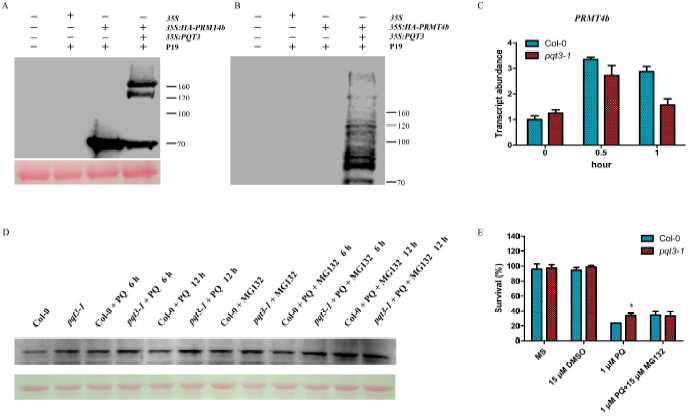
PRMT4b is a substrate recognized by PQT3. **(A and B)** *In planta* ubiquitination assay. Total protein samples were isolated from the infiltrated parts of *N. benthamiana* leaves 1 day after agroinfiltration. The total protein was analyzed via western blot using anti-HA antibody **(A)**. The total proteins were separated via a SDS-PAGE and Ponceau S staining of the Rubisco protein was shown as a loading control. Immunoprecipitated samples were analyzed using western blot with anti-ubiquitin antibody subsequently **(B)**. The numbers on the right indicate the molecular masses of marker proteins. **(C)** The mRNA level of *PRMT4b* in wild type and *pqt3* mutant under paraquat treatment. Total RNA was isolated from 14-day-old wild type and *pqt3* seedlings without or with 6 μM paraquat treatment for quantitative RT-PCR analysis. Values are mean ±SD (n=3 experiments). **(D)** The effect of proteasome inhibitor MG132 on the protein level of PRMT4b in wild type and *pqt3-1* mutant with paraquat treatment in time-course. Wild type and *pqt3-1* mutant were grown for 14 days, then the seedlings were treated without or with 6 μM paraquat for 6 or 12 h in presence or absence of 50 μM proteasome inhibitor MG132. The total proteins were extracted. Western blot was performed using anti-PRMT4b antibody. Ponceau S staining of the Rubisco protein serves as a loading control. **(E)** The survival ratio of wild type and *pqt3* mutant germinated and grown on MS medium containing 0 μM or 1 μM paraquat without or with 15 μM MG132 for 7 days were counted. DMSO was used as control. Values are mean ±SD (n= 30 plants, *P < 0.05). Asterisk indicates Student’s t-test significant difference.

The transcript level of *PRMT4b* could be induced in both wild type and *pqt3* seedlings by paraquat treatment (Figure 7C). The mRNA level of *PRMT4b* was similar between wild type and *pqt3* seedlings without or with paraquat treatment for 0.5h. The transcript level of *PRMT4b* in *pqt3* seedlings was lower than that of the wild type under paraquat treatment for 1h (Figure 7C). The PRMT4b protein levels were also detected using the seedlings treated without or with paraquat. As shown in the Figure 7D, the protein level of PRMT4b in *pqt3-1* was higher than that of the wild type under normal conditions. The protein level of PRMT4b in wild type remains lower than that of *pqt3-1* under paraquat treatment at the same time point, although protein level of PRMT4b increased gradually in wild type and *pqt3-1* with the prolonged paraquat treatment (Figure 7D). Under MG132 treatment, the protein level of PRMT4b was increased in both wild type and *pqt3-1* mutant. When the seedlings were co-treated with paraquat and MG132 for 12 h, PRMT4b was accumulated in both wild type and *pqt3-1* mutant, and no significant difference of PRMT4b protein level was found between the wild type and *pqt3-1* mutant (Figure 7D) because the degradation of the PRMT4b protein through ubiquitination-26S proteasome pathway was inhibited by MG132.

In order to further demonstrate the PQT3-dependent ubiquitination of PRMT4b, the phenotype of wild type and *pqt3-1* mutant treated without or with paraquat in presence or absence of proteasome inhibitor MG132 was studied. Survival ratio of *pqt3-1* seedlings was higher than that of wild type under paraquat treatment. No significant difference of survival could be observed when the wild type and *pqt3-1* seedlings were co-treated with paraquat and MG132 (Figure 7E). The wild type gained enhanced paraquat tolerance as *pqt3-1* mutant under MG132 treatment.

### *PRMT4b* is involved in the oxidative stress tolerance

To reveal whether PQT3-enhanced the oxidative tolerance of plants is via PRMT4b in Arabidopsis, we obtained the *prmt4b* mutant and *35Spro:PRMT4b* lines (Supplemental Figures 1G, 1H and 1L) and observed the phenotypes of *prmt4b* and *35Spro:PRMT4b* under different concentrations of paraquat treatment firstly. Survival ratio of wild type, *pqt3-1*, *pqt3-2*, *prmt4b* and *35Spro:PRMT4b* were counted (Figure 8A). *35Spro:PRMT4b* had similar phenotype as *pqt3-1* and *pqt3-2*, while the *prmt4b* mutant was more sensitive to paraquat treatment than wild type (Figure 8A). Under CdCl_2_ treatment, primary root elongation of *prmt4b* mutant was also slower than that of wild type (Figure 8B). Furthermore, the overexpression lines of *PRMT4b* were analyzed under other stress conditions. *35Spro:PRMT4b* increased tolerance to CdCl_2_ and NaCl stresses, as compared with wild type. The *35Spro:PRMT4b* showed the opposite of the *35Spro:PQT3* which was more sensitive to CdCl_2_ and NaCl stresses as compared with wild type (Supplemental Figure 5). These results show that PRMT4b is a positive regulator for plant oxidative tolerance, which is also consistent with the function of PQT3. The *prmt4a* mutant and *prmt4aprmt4b* double mutants were subsequently obtained and examined (Supplemental Figures 1I to 1K). The *prmt4aprmt4b* double mutants had the similar phenotype with *prmt4b* under paraquat treatment, and the knockout of *PRMT4a* did not affect the oxidative tolerance (Figures 8C to 8E).

**Figure 8.**
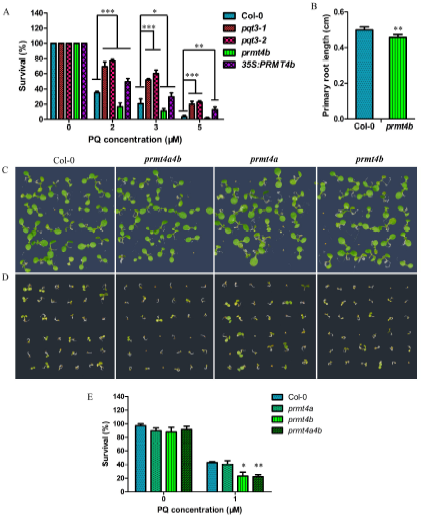
PRMT4b was involved in oxidative tolerance of Arabidopsis. **(A)** Paraquat tolerance assay. Survival ratio of wild type, *pqt3-1* mutant, *pqt3-2* mutant (Salk_065409), *prmt4b* mutant and *35Spro:PRMT4b* grown on 0, 2, 3, or 5 μM paraquat medium were counted. The assay was repeated for three times. Values are mean ±SD (n= 50 plants, *P < 0.05, **P < 0.01, ***P < 0.001). Asterisks indicate Student’s t-test significant differences. **(B)** Primary root elongation of wild type and *prmt4b* mutant seedlings grown on MS without or with 150 mM CdCl_2_ was measured. Values are mean ±SD (n= 30 plants, **P < 0.01). Asterisks indicate Student’s t-test significant differences. **(C to E)** The phenotype of wild type, *prmt4a*, *prmt4b*, and *prmt4aprmt4b* grown on MS without **(C)** or with 1 μM paraquat **(D)**. The survival ratio of wild type, *prmt4a*, *prmt4b*, and *prmt4aprmt4b* germinated and grown on MS medium containing 0 μM or 1 μM paraquat for 5 days were counted **(E)**. Values are mean ±SD (n= 36 plants, *P < 0.05, **P < 0.01). Asterisks indicate Student’s t-test significant differences.

### Increased H3R17 methylation on *APX1* and *GPX1* chromatin in *pqt3*

The transcript level of *APX1* and *GPX1* was higher in *pqt3* than that in wild type under normal conditions (Figures. 4A and 4B). PQT3 may regulate the transcription of *APX1* and *GPX1* via PRMT4b. Therefore, chromatin immunoprecipitation (ChIP) assays were carried out to compare the modification status of H3R17me2a in the chromatin of *APX1* and *GPX1* between wild type and *pqt3* mutant. ChIP assays were performed with wild type and *pqt3* plants using antibody against H3R17me2a. As shown in the Figures 9A and 9B, the *APX1* and *GPX1* chromatin was divided into different regions and the enriched chromosome fragments were detected by quantitative RT-PCR. The results showed that histone H3R17me2a modification of *APX1* and *GPX1* chromatin was increased in *pqt3* mutant (Figures 9C and 9D). These results suggest that PRMT4b may target *APX1* and *GPX1* to enhance the oxidative tolerance by increasing asymmetric dimethylation of H3 at R-17 in *APX1* and *GPX1* chromatin.

**Figure 9.**
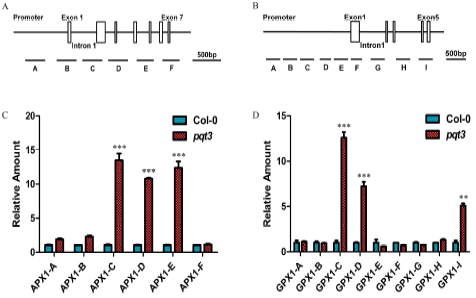
Arg-17 methylation in specific regions of *APX1* and *GPX1* chromatin was increased in *pqt3* mutant. **(A and B)** The illustration of *APX1* and *GPX1* chromatin. A to F represent different regions of *APX1* chromatin **(A)**. A to I represent different regions of *GPX1* chromatin **(B)**. Promoters (black lines), exons (white boxes), and introns (black lines) of *APX1* and *GPX1* were shown. **(C and D)** ChIP-PCR assay. Quantitative PCR was performed to verify each chromatin region of *APX1* and *GPX1* using specific primers. Fragments C, D and E in *APX1* chromatin were enriched by anti-H3R17 antibodies **(C)**. Fragments C, D and I in *GPX1* chromatin were enriched by anti-H3R17 antibodies **(D)**. *UBQ5* was used as an internal control. Values are mean ±SD (n=3 experiments, **P < 0.01, ***P < 0.001). Asterisks indicate Student’s t-test significant differences.

### Genetic analysis of PRMT4b acting as a target of PQT3 in oxidative tolerance

To confirm further that PRMT4b is the target of PQT3, the *pqt3prmt4b* double mutants were obtained (Supplemental Figures 1M and 1N). The survival ratio of *pqt3prmt4b* under paraquat treatment was intermediate between *pqt3* and *prmt4b*, which demonstrates that the PRMT4b protein is one of the targets of PQT3 and suggests that PQT3 may also target other proteins that contribute to the tolerance to oxidative stress (Figures 10A to 10C).

**Figure 10.**
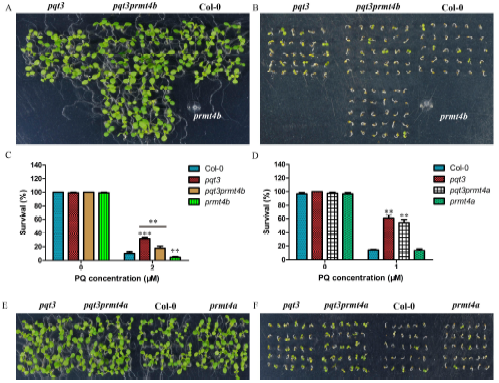
The phenotypes of *pqt3prmt4b* and *pqt3prmt4a* double mutants under paraquat treatment. **(A and B)** The phenotypes of *pqt3*, *pqt3prmt4b*, wild type and *prmt4b* seedlings grown on MS without **(A)** or with 2 μM paraquat **(B)**. **(C)** The survival ratio of *pqt3*, *pqt3prmt4b*, wild type and *prmt4b* under 2 μM paraquat treatment for 7 days were counted. Values are mean ±SD (n= 30 plants, **P < 0.01, ***P < 0.001). Asterisks indicate Student’s t-test significant differences. **(D)** The survival ratio of *pqt3*, *pqt3prmt4a*, wild type and *prmt4a* under 1 μM paraquat treatment for 7 days were counted. Values are mean ±SD (n= 30 plants, **P< 0.01). **(E and F)** The phenotypes of *pqt3*, *pqt3prmt4a*, wild type and *prmt4a* seedlings grown on MS without **(E)** or with 1 μM paraquat **(F)**.

We also obtained the *pqt3prmt4a* double mutants (Supplemental Figures 1O and 1P) and found that the survival ratio of *pqt3prmt4a* under paraquat treatment was not significantly different from that of *pqt3* mutant (Figures 10D to 10F), again indicating that PRMT4a is not involved in oxidative stress response.

## Discussion

PQT3 is a member of Arabidopsis RING-finger/U-box E3 ligase family. Secondary structure prediction using InterProScan protein sequence analysis software revealed four conserved domains including DWNN (domain with no name), zinc finger domain, RING-finger domain, and U-box domain in N-terminus of PQT3 protein (Supplemental Figures. 6A and 6B). DWNN is a novel ubiquitin-like domain, which is a highly-conserved domain in eukaryotic animals and plants (Pugh et al., 2006). The DWNN domain of PQT3 contains 76 amine acids. DWNN domain is only found in the N-terminus of the members in the splicing-associated RBBP6 (Retinoblastoma Binding Protein 6) protein family (Pugh et al., 2006). The RING-finger domain was also found in RBBP6 protein family. The existence of the RING-finger domain suggests that DWNN domain may act as ubiquitin-like regulatory factors (Pugh et al., 2006). CCHC-type Zinc finger domain is also known as zinc knuckle, which can be found in a large number of RNA binding proteins (Lopato et al., 1999; Krishna et al., 2003). As mentioned above, RING-finger domain can combine with the E2 in the cascade reaction of ubiquitination system,while U-box is a modified RING-finger (Freemont, 2000). In addition, two predicted NLS sequences (471-477 and 696-711 amino acids) were also found in the C-terminus of PQT3, which is consistent with nuclear localization of the protein (Figures 2H to 2P; Supplemental Figures 6A and 6B). Phylogenetic tree analysis further revealed the high homology proteins of PQT3 in other species (Supplemental Figure 6C). DWNN domain and RING-finger/U-box domain in N-terminus of PQT3 were highly conserved in homologous proteins in different plant species (Supplemental Figure 7). The PQT3 may play its role as an ubiquitin ligase in different species and its function may be conserved throughout the plant kingdom.

*In vitro* ubiquitination assay shows that PQT3 has the E3 ligase activity (Figure 5). For *in planta* ubiquitination assay, as compared with the control sample, the enhanced high molecular size bands in the PQT3-PRMT4b co-infiltration sample indicate that PRMT4b protein could be ubiquitinated by expressed PQT3 protein in *N. benthamiana* (Figures 7A and 7B). The transcript level of *PRMT4b* was not significantly different between wild type and *pqt3* mutant without or with paraquat treatment for 0.5 h (Figure 7C). The transcript level of *PRMT4b* was lower in *pqt3* mutant than that in wild type under paraquat treatment for 1 h (Figure 7C). One possible reason is that *pqt3* has enhanced oxidative tolerance to paraquat which could induce the transcript level of *PRMT4b* (Figures 1 and 7C). In addition, the reduced transcript level of *PRMT4b* in *pqt3* may be due to a negative feedback regulation that is activated by the accumulation of PRMT4b protein under paraquat treatment (Figure 7D). In Arabidopsis, *pqt3-1* and *pqt3-2* had higher protein level of PRMT4b than the wild type, as the PRMT4b protein was degraded by PQT3 in wild type under normal conditions (Figure 7D). The transcript level of *PRMT4b* could be induced in wild type under paraquat treatment (Figure 7C), while the transcript level of *PQT3* was decreased by paraquat treatment (Figure 3A). PQT3-mediated PRMT4b degradation was weakened in wild type. Consequently, the level of PRMT4b protein in wild type was elevated, but it was lower than that of *pqt3* (Figure 7D). Proteasome inhibitor MG132 could block the degradation of the PRMT4b protein in wild type and enhance paraquat tolerance of wild type (Figures 7D and 7E). Above all, PQT3, as an E3 ligase, play its role in oxidative tolerance through the ubiquitination-degradation of PRMT4b.

A series of environmental stress could lead to oxidative damage to plants (Zhu, 2000; Mittler, 2002; Apel and Hirt, 2004). Both biotic stress and abiotic stress result in the production of ROS which in excess cause oxidative stress. Besides paraquat tolerance, *pqt3* mutants have enhanced tolerance to various environment stresses (Figure 1; Supplemental Figure 2). Further studies may reveal new mechanisms of PQT3 in multiple stress tolerance. Analysis of PQT3-interacting proteins may be a good start point to further understand the function of *PQT3*. In the screen for PQT3-interacting proteins with Y2H, we isolated 47 colonies that were positive for the expressions of the His3 and LacZ reporter genes. Among these proteins, PRMT4b, as previously mentioned, may be responsible for the increased degree of Arg-17 methylation which may further regulate *APX1* and *GPX1* genes to enhance oxidative tolerance of plants (Figures 4A, 4B and 9). It has been reported that arginine methylation was involved in transcriptional regulation, signal transduction, DNA repair, RNA processing, and nuclear transport (Bedford and Richard, 2005; Pahlich et al., 2006; Bedford and Clarke, 2009; Liu et al., 2010a; Yu, 2011; Ahmad and Cao, 2012; Blackwell and Ceman, 2012). PRMTs have been extensively analyzed, and the functions of PRMTs involved in flowering time, circadian cycle, vegetative growth, and the plant response to salinity and ABA have been reported (Niu et al., 2008; Ahmad and Cao, 2012). Here, the *prmt4b* mutant was found to be more sensitive to paraquat and CdCl_2_ treatment than wild type (Figure 8). Thus, PRMT4b plays a role in plant response to oxidative stress. It is known that PRMT4-mediated methylation at Arg-17 of histone H3 is linked to transcription activation (Bauer et al., 2002). The Arg-17 of histone H3 is the major site of PRMT4-mediated methylation, although it was reported that other protein site could also be methylated by PRMT4 (Chevillard-Briet et al., 2002; Fujiwara et al., 2006; Cheng et al., 2007). We suggest that the increased Arg methylation degree of specific regions in *APX1* and *GPX1* chromatin was caused by PRMT4b (Figure 9).

The increased transcript level of *APX1* and *GPX1* may be caused by PRMT4b-mediated histone Arg methylation (Figures 4A, 4B and 9). APX1 is a cytosolic enzyme involved in H_2_O_2_ degradation using ascorbate as the electron donor (Correa-Aragunde et al., 2013). The response of *APX1* to oxidative stress has been studies in Arabidopsis. APX1 plays an important role in the response to stress combination (Koussevitzky et al., 2008). It could be activated by multiple stresses to protect plants against oxidative stress (Davletova et al., 2005; Vanderauwera et al., 2011; Maruta et al., 2012). The tobacco could be more tolerant against UV-C-induced oxidative damage through the overexpression of *APX1* (Saxena et al., 2011). GPXs also have important functions in oxidative signaling, which can protect plants from harmful effects of excessive oxidation (Passaia et al., 2014). It has also been reported that overexpression of a tobacco glutathione S-transferase with glutathione peroxidase activity (GST/GPX) in transgenic tobacco seedlings lead to enhanced peroxide scavenging and reduced oxidative damage (Roxas et al., 2000).

The *pqt3* mutants also have late-flowering phenotype (Figure 1D). It has been reported that *prmt4aprmt4b* double mutants display late-flowering phenotype (Niu et al., 2008). The Y2H result showed that the PQT3 can not interact with PRMT4a (Supplemental Figure 4). As compared with wild type, the *prmt4a* mutant also has no significant difference in the oxidative tolerance (Figures 8C to 8E). The phenotype of *pqt3prmt4a* mutant demonstrates that PRMT4a protein was not involved in the regulation of oxidative stress response by PQT3 (Figures 10D to 10F). The late-flowering phenotype of *pqt3* may be regulated by other mechanisms, rather than *PRMT4a* and *PRMT4b*. The flowering-related transcription factor AGAMOUS (AG) was found to be a potential target interacted with PQT3 in Y2H library screening. AG is involved in carpel development, leaf development, identification of floral organs, and stamen development (Urbanus et al., 2010). Targeted removal of AG by PQT3 may be related to the late-flowering phenotype of *pqt3*. By interacting with different partners, PQT3 may mediate multiple functions in diverse biological processes.

In conclusion, oxidative stress activates the expression of *PRMT4b*, represses the expression of *PQT3*, and weakens PQT3-mediated the ubiquitinated degradation of PRMT4b, synergistically resulting in increased accumulation of PRMT4b. Consequently, the increased level of PRMT4b protein may lead to higher degree of histone methylation on the *APX1* and *GPX1* chromatin. As a result, the transcription of *APX1* and *GPX1* is activated, leading to more APX1 and GPX1, which enhance oxidative tolerance of plants. When the stress disappears, transcription repression of *PQT3* by oxidative stress is removed. The function of PQT3, as a negative regulator of oxidative stress response, is restored. The PRMT4b is then degraded by PQT3 in ubiquitination pathway. The activated response to oxidative stress is switched off. We propose a working model for PQT3 as a negative regulator of oxidative stress response (Figure 11).

**Figure 11.**
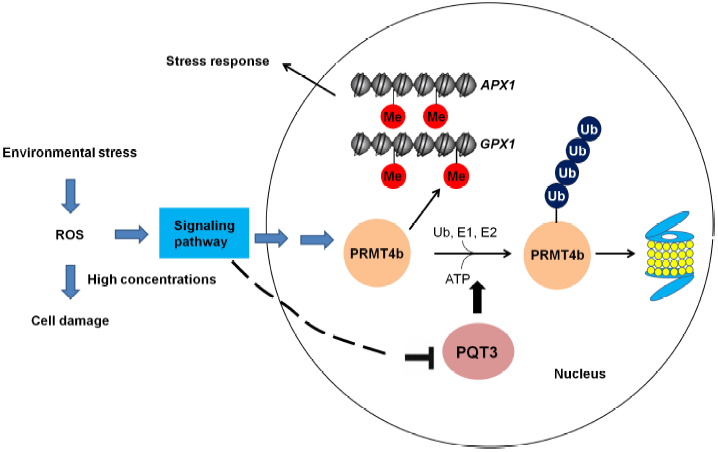
A working model for PQT3 acting as a negative regulator of oxidative stress response. Many environmental stresses cause oxidative stress in plants. Under oxidative stress, stress signaling up-regulates *PRMT4b* expression and down-regulates *PQT3* expression, leading to higher PRMT4b activity that will activate *APX1* and *GPX1* and enhance antioxidation capacity of APX1 and GPX1. When oxidative stress is diminished, *PRMT4b* expression is decreased and *PQT3* expression is increased. As a result, PQT3 activity is increased, leading to faster removal of PRMT4b via 26S proteasome. Together with decreased *PRMT4b* expression, PRMT4b activity will rapidly drop, leading to decreased expression of *APX1* and *GPX1*. The activated response of oxidative stress is then switched off.

## Methods

### Mutant screen from the activation-tagging library

The paraquat tolerant mutant *PQT3* was isolated from an activation-tagging library including approximately 55,000 individual lines generated using pSKI015 in the Columbia ecotype as described (Xi et al., 2012).

### Plant material and growth conditions

The wild type used in the study is *Arabidopsis thaliana* ecotype Columbia that is the genetic background for all the mutants and transgenic plants. Salk_065409 (*pqt3-2*), Salk_097442C (*prmt4b*) and Salk_033423 (*prmt4a*) were ordered from ABRC (Arabidopsis Biological Resource Center). The seeds were sterilized in 10% bleach for 10 min. Then the seeds were washed for 5 times at least with sterile water. For vernalization, the seeds were kept in the dark with water at 4°C for 3 days to ensure the synchronous germination. Sterile seeds were germinated on half-strength MS medium. The seedlings were grown at 22°C under 16-h-light (100 μE m^−2^ s^−1^) /8-h-dark cycle.

### Transformation of Arabidopsis

The constructs were electroporated into competent cell of *Agrobacterium tumefaciens* C58C1. The floral-dip method was used to transfer these constructs into Arabidopsis as described (Clough and Bent, 1998; Bent, 2000).

### PCR analysis

Total RNA was extracted from various tissues of plants using TRIzol reagent. Then RNA reverse reaction was carried out by TransScript Kit (TransGen Biotech). For RT-PCR analysis, PCR was performed using specific primers and the products were detected by agarose gel electrophoresis. For quantitative RT-PCR analysis, the transcript level was detected on an Applied Biosystem Step One real-time PCR system using SYBR Premix Ex Taq II (TaKaRa) and specific primers were listed in the Supplemental Table 1. *UBQ5* was used as the internal control.

### Identification of *pqt3* mutant, FC line, *35Spro:PQT3*, and *35Spro:PRMT4b*

Homozygous T-DNA insertion mutants of Salk_150614 were identified using genomic PCR as described (Alonso et al., 2003). RT-PCR was carried out as previously described to confirm the results of genomic PCR. The *35Spro:PQT3* and *35Spro:PRMT4b* plasmids were transformed into Col-0 to obtain the overexpression lines of *PQT3* and *PRMT4b*. For the FC line, the *35Spro: PQT3* was transformed into *pqt3* mutant, and the line with the same expression level of *PQT3* as wild type was chosen and used as FC line. *35Spro: PQT3, 35Spro: PRMT4b* and FC line were identified by glufosinate screening and RT-PCR.

### Stress tolerance assay

The seeds were germinated on half-strength MS media containing different concentrations of paraquat, mannitol, CdCl_2_ and NaCl, respectively. The phenotype was observed and survival ratio was scored at the indicated time points.

For drought tolerance assay, the *pqt3-1* mutant, *pqt3-2* mutant and wild type seeds were germinated in one pot at same density. When seedlings were 15 day old, watering was withheld for 15 day before re-watering. The photos were taken before re-watering and after re-watering for 1 day and 7 days. The survival ratio was scored after re-watering for 1 day and 7 days.

### DAB staining

DAB staining was performed as described (Daudi and O’Brien, 2012). DAB staining solution (pH 6.0) was prepared by adding 0.05% (v/v) Tween-20 and 10 mM Na_2_HPO_4_ to the DAB solution (1 mg/ml DAB, pH3.0). For each treatment condition, at least 3 leaves per plant were obtained from 3 independent plants for each line (Col-0, *pqt3-1* and *pqt3-2*). Arabidopsis leaves from different lines were treated using MS liquid medium without or with 6 μM paraquat for 3 h. These leaves were stained in 6-well culture plates with DAB staining solution subsequently and 10 mM Na_2_HPO_4_ (pH 6.0) was used as the negative control. The 6-well plates were covered with alumium foil and placed on a shaker for 4 - 5 h. Follow the incubation, replace the DAB staining solution with bleaching solution (ethanol: acetic acid: glycerol = 3:1:1). Place the 6-well plates into a boiling water bath (90 - 95°C) for 15 - 20 mins. Replace the bleaching solution with fresh bleaching solution. The brown precipitate formed by the DAB reaction with H_2_O_2_ could be observed on the leaves. Photos were taken using a camera. Special attention should be paid to light avoidance through the whole operation. The experiment was repeated for three times.

### The detection of GUS activity

The promoter of *PQT3* was cloned into pCB308R (Xiang et al., 1999; Lei et al., 2007). The transgenic lines containing *PQT3pro: GUS* were isolated by glufosinate screening. The T2 population was used for GUS staining. Histochemical staining for GUS activity in Arabidopsis was performed as described previously (Jefferson et al., 1987) and GUS staining solution was prepared as described before (Xi et al., 2012).

The experimental materials were incubated in staining solution at 37°C. Then Arabidopsis tissues were destained and stored in 70% ethanol. The GUS activities of individual parts were observed via a light microscope (ZEISS Axio skop2 plus) with a video camera.

### Subcellular localization assay

The full length CDS of *PQT3* was cloned into the binary vector pCB2008E to construct the PQT3-GFP fusion vector (Lei et al., 2007). The inserted sequence was confirmed by sequencing. The construct was delivered into the onion epidermal cells via microprojectile bombardment of particle gun for transient expression assay. It was also transferred into Col-0 via floral-dip method to create transgenic plant for the analysis of PQT3-GFP fusion protein localization. The green fluorescence in onion epidermal cells and the root tissues of transgenic lines were observed using a fluorescence microscope (ZEISS Axio skop2 plus) with a video camera.

### Enzyme activity assay

For APX and GPX enzyme activity assay, Arabidopsis seedlings were ground in liquid nitrogen and resuspended in precooling Enzyme extraction buffer (50 mM phosphate buffer (Na_2_HPO_4_-NaH_2_PO_4_), PH=7.0; 1 mM EDTA; 0.05% (v/v) TritonX-100; 2% (w/v) PVPP and 1 mM ascorbic acid) on ice. Extraction solution was centrifuged for 20 min (16000 g/min, 4°C). The protein concentration of supernatant was detected by One-drop micro-ultraviolet spectrophotometer and SDS-PAGE before the supernatant was used for enzyme activity analysis. The detection of APX activity was performed as described with modifications (Nakano and Asada, 1981). For APX activity, 50 μl enzyme and 2950 μl rection mixture (50 mM Tris-HCl, pH7.0; 0.1 mM EDTA; 0.1 mM H_2_O_2_ and 0.5 mM ascorbic acid) was mixed. The decreased OD290 was recorded per 10 s. The amount of enzyme oxidized 1 mM AsA in one minute set as one activity unit (U) of APX. The activity of APX was expressed as U • g-^1^ protein. GPX activity was measured indirectly through the detection of glutathione reductase (GR) activity using GPX Activity Measurement Kit (Beyotime Biotech, China). GR activity was detected as described with modifications (Halliwell and Foyer, 1978). The OD340 was recorded per 30 s. The amount of enzyme consumed 1 mM NADPH in one minute set as one activity unit (U) of GPX. The activity of GPX was also expressed as U • g-^1^ protein.

### Western blot

For western blot analysis, proteins were separated by SDS-PAGE in a 12% acrylamide gel and electroblotted to nitrocellulose membrane (Immobilon-P, MILLIPORE Corporation, USA). Antibodies used in western blot were as follows: anti-HA antibody (HA-Tag, 26D11, Mouse mAb, M20003, Abmart, Shanghai, China), 1:1000 for western blot; anti-PRMT4b antibody, 1:500 for western blot; anti-Ubiquitin antibody (ab7254, abcam, USA), 1:1000 for western blot; anti-His antibody (His-Tag, 2A8, Mouse mAb, M30111, Abmart, Shanghai, China), 1:1000 for western blot and goat anti-mouse lgG-HRP (Santa Cruz Biotechnology, USA), 1:5000 for western blot. The results were examined with a CCD camera system (Image Quant LAS 4000) using Super Signal West Femto Trial Kit (Thermo, USA).

### *In vitro* E3 ubiquitin ligase activity assay

The *in vitro* E3 ligase activity assay was performed as described previously (Zhang et al., 2007). GST-PQT3 fusion protein was obtained from *E. coil* and purified subsequently. His-tagged Arabidopsis ubiquitin (UBQ14) was also expressed using bacterial expression system and purified. In addition, the wheat *(Triticum aestivum)* E1 (GI: 136632) and human E2 (UBCh5b) were also used in the reaction. Reactions were performed for 1.5 h at 30°C. For the immunoblot, Nickel-HRP (nickel–nitrilotriacetic acid agarose conjugated to horseradish peroxidase) (Kirkegaard & Perry Laboratories, http://www.kpl.com/) was used for the detection of His-tagged ubiquitin.

### Y2H screening and confirmation

The full-length *PQT3* cDNA was cloned into the bait vector pDEST32 and transformed into the yeast strain Mav203 subsequently. Yeast two-hybrid screening was performed using two-hybrid cDNA library of Arabidopsis. Yeast cells harboring the bait were transformed with a cDNA library containing inserts for prey proteins fused to GAL4-AD. Positive clones were screened using SD/-Leu-Trp-His and X-gal assay. Then it was identified by nucleotide sequencing with corresponding primers. Two-hybrid screening was performed via the protocol described in Two-Hybrid System Manual (Invitrogen, USA). The yeast two-hybrid was carried out to confirm the result of screening. The full-length CDS of *PQT3* and the four segments of *PQT3* were cloned into the bait vector pDEST32 and *PRMT4b* was cloned into the prey vector pDEST22. The primers used for yeast two-hybrid were listed in Supplemental Table 1.

### Pull-down assay

MBP-PQT3-C66 and His-AtPRMT4b fusion protein were expressed using prokaryotic expression system and purified. MBP-PQT3-C66 fusion protein was incubated with MBP beads (amylose resin) at 4°C for 2 h, and the MBP tag was used as a negative control. The beads were cleaned with washing buffer for 4 times. Then the beads were incubated with His-AtPRMT4b at 4°C for 2 h respectively. The beads were cleaned with washing buffer for 4 times. Pulled-down mixtures were separated by SDS-PAGE and further transferred to nitrocellulose membrane. The result was detected using western blot with anti-His antibody.

### Agroinfiltration procedure

*Agrobacterium tumefaciens* strain C58C1 was used in the experiments. Agroinfiltration procedure was performed as described previously (Liu et al., 2010b). At first, these strains were grown on LB medium containing Kanamycin. Single colony was transferred into 5 ml LB liquid medium with Kanamycin and grown for 48 h in a 28°C shaker. The bacteria solution was inoculated into new LB liquid medium containing 10 mM 2-(N-morpholine)-ethanesulfonic acid (MES; pH 5.6) and 40 μM acetosyringone (1:100 ratio, v/v). Bacteria were developed in a 28°C shaker until OD_600_ reached 3.0 approximately. The bacteria were collectted gently by means of 10 min centrifugation (3200 g/min), and the pellets were resuspended with 10 mM MgCl_2_ until OD_600_ reached 1.5 approximately. The bacteria solution was kept at room temperature with a final concentration of 200 μM acetosyringone for at least 3 h without shaking. The different plastid combinations were transformed into epidermal cells of *N. benthamiana* leaves by disposable syringe.

### BiFC analysis

NE-PQT3 (the N-terminal part of YFP fused with PQT3) and CE-PRMT4b (the C-terminal part of YFP fused with PRMT4b) were constructed. These constructs were transferred to Agrobacterium strains C58C1 respectively. As mentioned above, the different plastid combinations were transformed into epidermal cells of *N. benthamiana* leaves by agroinfiltration. YFP was observed 1-2 days after leaf infiltration using confocal. The nuclei were stained by Hoechst subsequently and fluorescence detection by confocal was performed. The primers used for BiFC were listed in Supplemental Table 1.

### Protein extraction

Native extraction buffer 1 [ NB1; 50 mM TRIS-MES pH 8.0, 0.5 M sucrose, 1 mM MgCl_2_,10 mM EDTA, 5 mM DTT, protease inhibitor cocktail for plant cell and tissue extracts (Sigma, USA) ] was chosen for protein extraction buffer. Other steps of protein extraction were carried out as described previously (Liu et al., 2010b).

### *In planta* ubiquitination assay

Agrobacterium strains containing different constructs were mixed and infiltrated into the leaves of *N. benthamiana*. Total proteins were extracted from plant leaves 1 day after infiltration. The total proteins were separated by SDS-PAGE and analyzed via western blot using anti-HA antibody. Ponceau S staining of the Rubisco protein was used as loading control. In the presence of MG132, total proteins were immunoprecipitated with anti-HA antibody (Abmart) and protein A agarose beads (Millipore, USA) subsequently. Detailed steps of immunoprecipitation were performed as described previously (Liu et al., 2010b). Immunoprecipitated samples were analyzed using western blot with anti-ubiquitin antibody (Abcam). The results were examined with a CCD camera system (Image Quant LAS 4000).

### Chromatin immunoprecipitation-PCR assay

The *pqt3* mutant and wild type were used for ChIP assay. *UBQ5* was used as an internal control. ChIP was performed as previously described (Gendrel et al., 2005; Pei et al., 2007). The regions with enriched Arg-17 methylation were precipitated from input DNA with anti-H3R17 antibodies (anti-Histone H3 asymmetric dimethyl R17 antibody-ChIP grade, ab8284, Abcam, USA). The enrichments of DNA fragments were detected by quantitative RT-PCR using specific primers corresponding to different regions of *APX1* and *GPX1* chromatins as described previously (Bastow et al., 2004; Liu et al., 2007). The primers used in ChIP assay were listed in Supplemental Table 1.

### Accession Numbers

Sequence data from this article can be found in the Arabidopsis Genome Initiative or GenBank/EMBL databases under the following accession numbers: *TUB8*, At5g23860; *UBQ5*, At3g62250; *PQT3*,At4g17410; *PRMT4b*, At3g06930; *PRMT4a*, At5g49020; *APX1*, At1g07890; *APX2*, At3g09640; *APX3*, At4g35000; *APX4*, At4g09010; *APX5*, At4g35970; *APX6*, At4g32320; *sAPX*, At4g08390; *tAPX*, At1g77490; *CSD1*, At1g08830; *CSD2*, At2g28190; *CAT1*, At1g20630; *CAT2*, At4g35090; *CAT3*, At1g20620; *FSD1*, At4g25100; *FSD2*, At5g51100; *FSD3*, At5g23310; *GPX1*, At2g25080; *GPX2*, At2g31570; *GPX3*, At2g43350; *GPX4*, At2g48150; *GPX5*, At3g63080; *GPX6*, At4g11600; *GPX7*, At4g31870; *GPX8*, At1g63460; *ACHT1*, At4g26160; *ACHT2*, At4g29670; *ACHT3*, At2g33270; *ACHT4*, At1g08570; *ACHT5*, At5g61440; *2CPB*, At5g06290; *PRXQ*, At3g26060; *GRXC1*, At5g63030; *GRXC2*, At5g40370; *GRXC5*, At4g28730; *MSD*, At3g10920; *AG*, At4g18960.

### Supplemental Data

Supplemental Figure 1. Identification of mutants, homozygous Salk line and transgenic lines.

Supplemental Figure 2. Phenotype of *pqt3* mutants under other environmental stresses leading to oxidative damage.

Supplemental Figure 3. Sections of PQT3 protein used in Y2H and pull-down assays.

Supplemental Figure 4. Y2H assay for PQT3 and PRMT4a.

Supplemental Figure 5. Phenotype of *PQT3* and *PRMT4b* overexpression lines under other environmental stresses lead to oxidative damage.

Supplemental Figure 6. Functional domains and phylogenetic tree of PQT3 protein.

Supplemental Figure 7. Homologous sequences alignment of PQT3 protein.

## ACKNOWLEDGEMENTS

This work was supported by National Natural Science Foundation of China (grant numbers 31270302 and 30770189 to C. X. and 31330020 to X. C.) and the National Basic Research Program of China (2011CB915400 to Q. X. and 2012CB114300 to C.X.). The authors thank the ABRC for providing the seeds of Salk_065409, Salk_097442C and Salk_033423, Dr. Lijia Qu (School of Life Sciences, Peking University, Beijing, China) for providing Arabidopsis cDNA library for Y2H screen, and Dr. Yong Ding (School of Life Sciences, University of Science and Technology of China, Hefei, China) for his critical review of the manuscript.

## AUTHOR CONTRIBUTIONS

C.L. and J.D. designed the study, performed the experiments and interpreted the results. XT.C. designed the study, performed the experiments, interpreted the results, and wrote the manuscript. P.Z. and R.L. contributed to part of the experiments. T.Z. and XF.C. contributed *prmt4b* mutants, *PRMT4b* overexpression lines, and PRMT4b antibody. P.W. and Q.X. contributed to the verification of E3 ligase activity of PQT3. C.X. designed the experiments, discussed the results, edited the manuscript and supervised the project.

